# Generating variability from motor primitives during infant locomotor development

**DOI:** 10.1101/2022.05.05.490063

**Authors:** Elodie Hinnekens, Marianne Barbu-Roth, Manh-Cuong Do, Bastien Berret, Caroline Teulier

**Affiliations:** Université Paris-Saclay, CIAMS, 91405, Orsay, France; Université d’Orléans, CIAMS, 45067, Orléans, France; Université de Paris, CNRS, Integrative Neuroscience and Cognition Center, 75006, Paris, France; Institut Universitaire de France, 75005 Paris, France

## Abstract

Motor variability is a fundamental feature of developing systems allowing motor exploration and learning. In human infants, leg movements involve a small number of basic coordination patterns called locomotor primitives, but whether and when motor variability could emerge from these primitives remains unknown. Here we longitudinally followed 10 neonates (∼4 days old) until walking onset (∼14 months old) and recorded the activity of their leg muscles during locomotor or rhythmic movements. Using unsupervised machine learning, we show that the structure of trial-to-trial variability changes during early development. In the neonatal period, infants own a minimal number of motor primitives but generate a maximal motor variability across trials thanks to variable activations of these primitives. A few months later, toddlers generate significantly less variability despite the existence of more primitives, due to more regularity within their activation. These results suggest that human neonates initiate motor exploration as soon as birth by variably activating a few basic locomotor primitives that later fraction and become more consistently activated by the motor system.

## Introduction

Despite its apparent simplicity when performed by healthy adults, walking is a complex task that involves numerous nerves, joints and muscles (Bernstein, 1967). To simplify the coordination of these numerous degrees of freedom (DOFs), the central nervous system (CNS) seems to efficiently control walking and other behaviors via a small number of encoded primitives, also called motor modules or muscle synergies (Bizzi et al., 1991; d’Avella et al., 2003; Tresch et al., 1999). A primitive is a neural structure that is stored within the CNS at a spinal level and that autonomously produces a coordinated pattern of behavior (i.e. involving several muscles) when recruited from higher centers (Bizzi et al., 2008; Overduin et al., 2012; Roh et al., 2011). Two types of primitives are described, including spatial modules and temporal modules. A spatial module is a group of muscles that are activated together with relative weights, while a temporal module is a waveform that describes the activation of a spatial module across time (Delis et al., 2014). In human, computational modeling from electromyographic (EMG) data led to the identification of 4 to 5 spatial and temporal modules involved in adult walking (Hinnekens et al., 2020; Ivanenko et al., 2004; Lacquaniti et al., 2012) (Figure 1).

**Figure 1.**
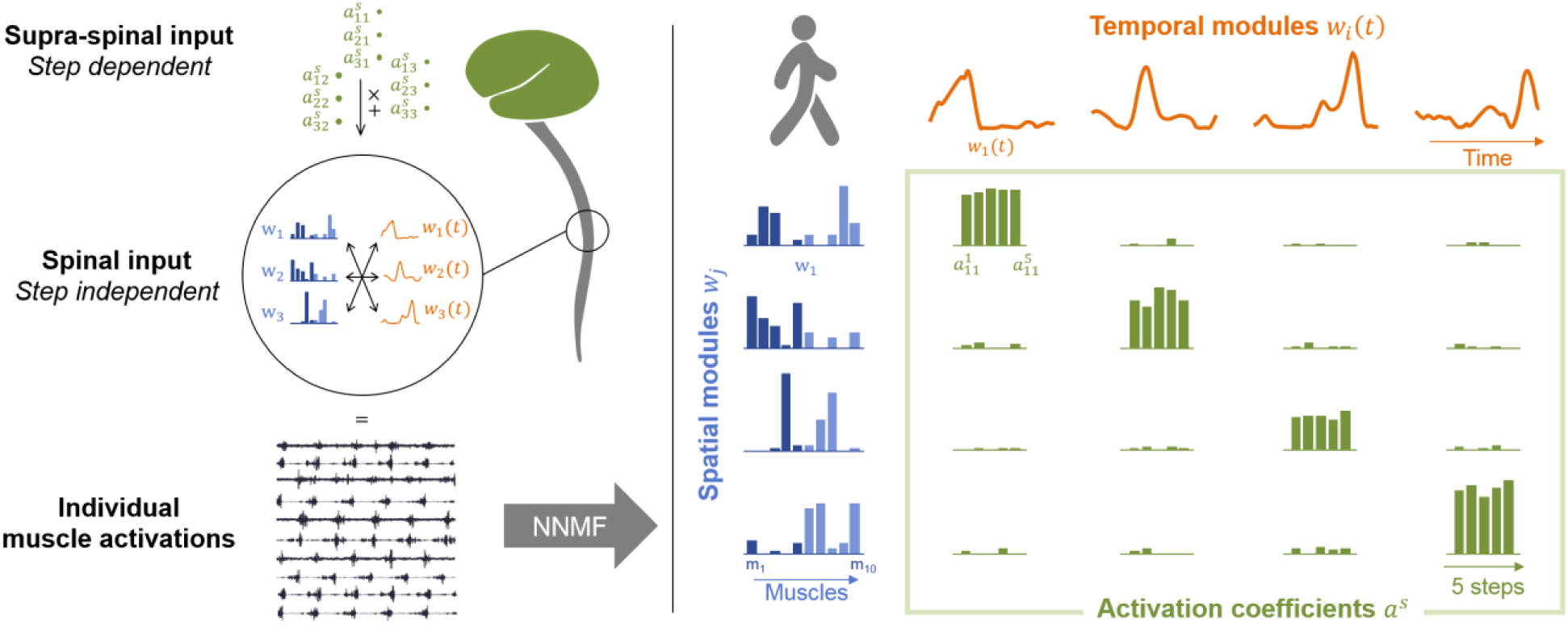
Theory of modularity and modular organization of adult walking. Left. The theory of modularity postulates that individual muscle activations result from the combination of basic spinal structures called locomotor primitives which are of two types: spatial (blue) and temporal (orange) modules. According to the space-by-time model that is used here, the brain activates those modules through a supraspinal input (green) that specifies which amplitude of activation has to be allocated to each possible pair of spatial and temporal modules. In humans, non-negative matrix factorization (NNMF) is used to identify the underlying motor primitives and their activation coefficients from EMG data. **Right**. Illustration of NNMF applied to 5 steps of walking in a human adult (Table S6). EMG patterns can be decomposed into 4 spatial modules (blue) and 4 temporal modules (orange). Within each spatial module, weightings are plotted for muscles m1 to m10 in the following order: rectus femoris, tibialis anterior, biceps femoris, soleus, and gluteus medius (right muscles in dark blue followed by left muscles in light blue). Activation coefficients (green) represent the level of activation of each possible pair of spatial and temporal modules during 5 steps. Two features are typical of adults’ modular organization: the stability of activation (activations coefficients remain stable during the 5 steps) and the selectivity of activation (one spatial module is always activated with only one temporal module and vice versa).

The development of this fine-tuned modular control has been investigated from birth on. Human locomotor precursors such as stepping or kicking have been found to already rely on a low-dimensional modular organization. For example, neonatal stepping is based on only 2 primitives while walking in toddlers is based on 4 primitives (Dominici et al., 2011; Sylos-labini et al., 2020). This is coherent with the fact that innate behaviors are described as stereotyped (Jeng et al., 2002; Spencer and Thelen, 2000; Thelen et al., 1981), with strong coupling among joints (Fetters et al., 2004; Jeng et al., 2002; Thelen et al., 1981) and among agonist/antagonist muscles (Teulier et al., 2012). However, while those observations suggest that neonates own a restricted motor repertoire, studies from behavioral literature report high intra-individual variability within neonatal movements. The muscular activity is notably highly variable during the first year of life (Teulier et al., 2012), which is believed to be a key feature of typical motor development allowing learning (Dhawale et al., 2017; Hadders-algra, 2018). Variability is indeed thought to be centrally regulated for the purpose of motor exploration (Kao et al., 2008; Mandelblat-Cerf et al., 2009) and is even described as a marker of typical development in humans (Hadders-algra, 2018).

If modularity and variability seem antagonistic at first glance, the question of whether those two features are compatible or incompatible needs to be addressed to better understand motor development. On one hand, a low-dimensional modular system inherently limits motor exploration (Cohn et al., 2018; Valero-Cuevas, 2009). In this vein the high variability of EMG data during the first year of life opened discussion about the existence itself of motor primitives (Teulier et al., 2012). On the other hand, data from animals suggest that variability could be generated within a modular system during development. A variable output was indeed observed after applying different stimuli to the neonatal spinal cord of rodents (Kiehn and Kjærulff, 1996; Klein et al., 2010), which is also believed to store motor primitives (Blumberg et al., 2013; Dominici et al., 2011). In young songbirds, a specialized cortical area has even been found as responsible for inserting variability into the temporal structure of vocalization to facilitate learning in early development, resulting in a highly variable output that becomes structured when inhibiting the area (Aronov et al., 2011; Kao et al., 2008). As those data suggest that is possible to produce variability by modulating the activation of basic inputs, such organization might shape the development of the motor system in human infants.

In human infants, investigations of the motor system are more limited and EMG recordings are the closest signals to the neural output that can be recorded while moving. Yet, if modularity and variability do coexist within the neural command, we should be able to separate the contribution of motor primitives from the variability of EMG signals and observe their cross evolution during development. To test this prediction, we longitudinally followed 10 human infants from birth to walking onset and recorded the EMG activity of 10 lower-limb muscles during leg movements at each age (Figure 2). Using a state-of-the-art unsupervised machine learning approach, we were able to model the underlying command by decomposing the EMG data of numerous muscles into step-invariant basic muscle patterns (which represent the motor primitives at each age) and into step-variable activation coefficients (which theoretically represent the variable descending command that modulates the activation of the motor primitives, at least in adults) (d’Avella et al., 2003; Delis et al., 2014) (Figure 1). We describe the evolution of both motor variability and motor modularity from birth to independent walking and provide evidence that the human motor system could theoretically initiate its exploration by variably activating a few temporary basic structured patterns.

**Figure 2.**
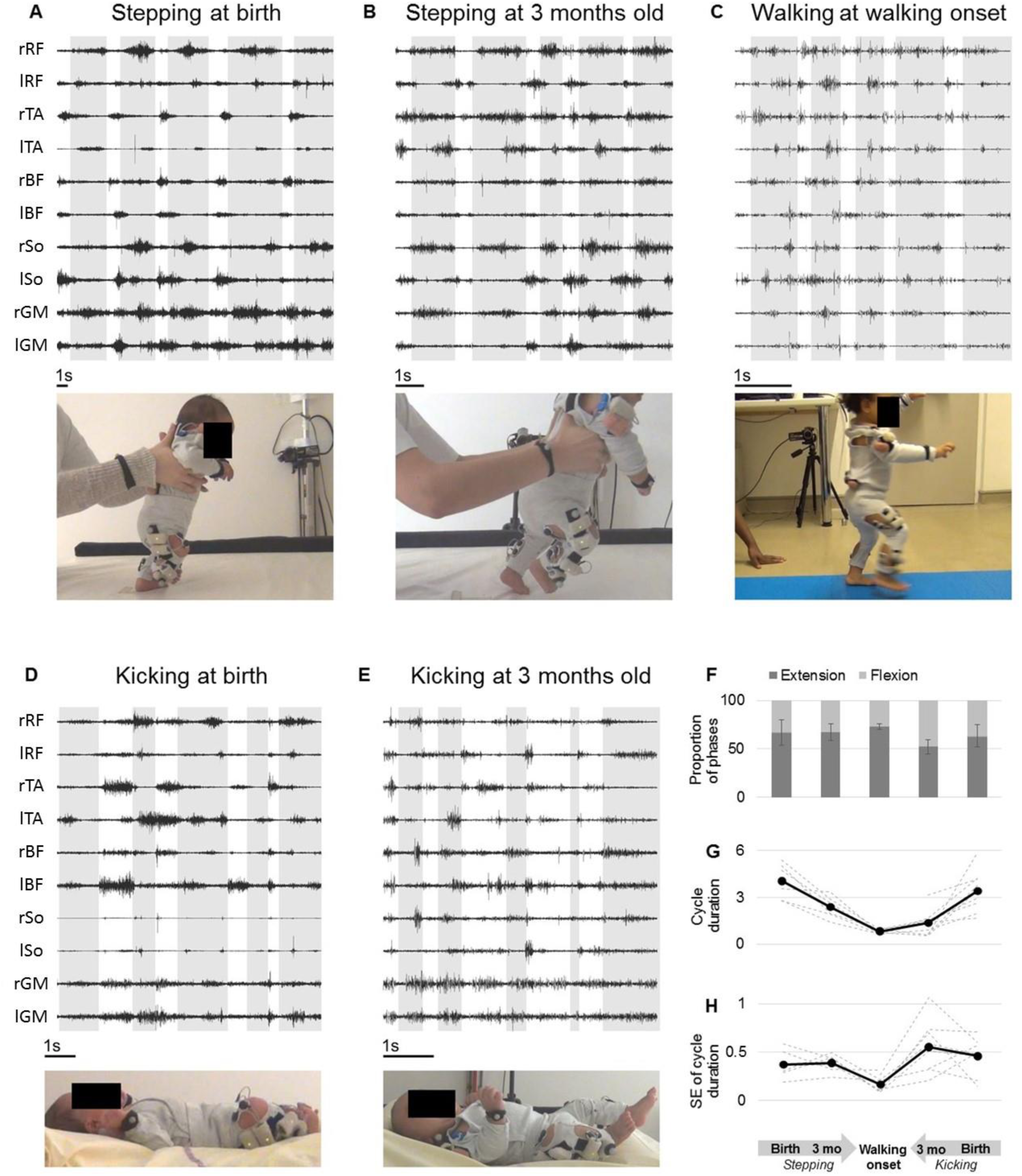
Development of basic EMG and kinematic parameters. **A to E:** example of EMG data for each age and behavior in one infant. Only the high-pass filter was applied in this representation. The scale of 1 second is display at the bottom of each figure. Extension phases appear on a grey background. RF Rectus Femoris, TA Tibialis Anterior, BF Biceps Femoris, So Soleus, GM Gluteus Medius **F to I:** Evolution of several features starting from birth to walking onset starting from either stepping or kicking. **F:** Proportion of flexion and extension phases. **G:** Cycle duration. **H:** Kinematic variability (standard deviation of cycle duration divided by averaged cycle duration).

## Results

12 infants were tested longitudinally from birth (∼4 days) to 3 months-old and 10 of them were tested shortly after they could walk independently (∼14 months). At birth and at 3 months, we observed the stepping behavior and/or the kicking behavior, while at walking onset we only recorded independent walking. In each behavior and at each age, infant’s movements were recorded using surface EMG on 10 bilateral lower-limb muscles and two 2D video cameras. Based on the resulting films, trained coders selected alternated cycles of flexion and extension of the lower-limbs, which allowed to study the same movement regardless of the behavior that could be produced by the infant at each age and to focus only on the generation of trial-to-trial variability for this given movement. Data from a given baby were considered analyzable when we had recorded clean surface EMG signals of the 10 lower-limb muscles during at least 5 alternated cycles of flexion and extension, both at birth and at 3 months old and through the same behavior (stepping or kicking). In total, 200 cycles of flexion and extension were included into the analysis (from 6 infants for stepping at birth and at 3 months old, 9 for kicking at birth and at 3 months, and 10 for walking; 5 cycles per behavior and per age for each infant). Then, for each behavior, we computed the variability of the motor output (index of EMG variability) and we used non-negative matrix factorization (NNMF) to identify the underlying motor primitives and their activation parameters. We computed a goodness-of-fit criterion to establish whether the trial-to-trial variability of 5 cycles of flexion and extension of the lower-limbs could be produced through various combinations of those motor primitives. We compared this goodness-of-fit criterion across ages and computed other indexes in order to characterize i) how variably were those motor primitives activated and ii) how selective were those primitives (i.e. if they controlled numerous muscles at a time or a few muscles). Procedure are detailed in the method section.

### Kinematic parameters and EMG signals reveal maximal motor variability during the neonatal period

We started by characterizing basic kinematic and EMG parameters at each age and in each behavior (Table S2). Wilcoxon tests were performed among kinematic parameters (cycle duration and its variability, proportion of extension/flexion phases) to assess basic differences. The cycle duration was different across behaviors with a decrease from stepping at birth to stepping at 3 mo (p=.03) and to walking in toddlers (p=.03) as well as a decrease from kicking at birth to kicking at 3 mo (p<.01) (Figure 2G). The proportion of phases within a cycle did not differ significantly across ages regarding stepping but some differences were observed regarding kicking (Table S3, Figure 2F).

The kinematic variability was assessed by the variability of cycle duration (Figure 2H). This variability significantly decreased for stepping and kicking from 3 mo to walking in toddlers (respectively p=.03 and p=.02). A significant decrease was also found from kicking at birth to walking (p=.02) as well as a trend toward significance from stepping at birth to walking (p=.06). However this variability did not seem to evolve between birth and 3 mo for stepping (p=.69) or kicking (p=.30). Regarding EMG data, the variability was assessed by the Index of EMG Variability (IEV, Figure 3D). This index significantly decreased with age in both stepping and kicking from birth to walking onset (p=.03 in both cases), as well as from 3 mo to walking onset for kicking (p=.047) and with a trend for stepping (p=.063).

**Figure 3.**
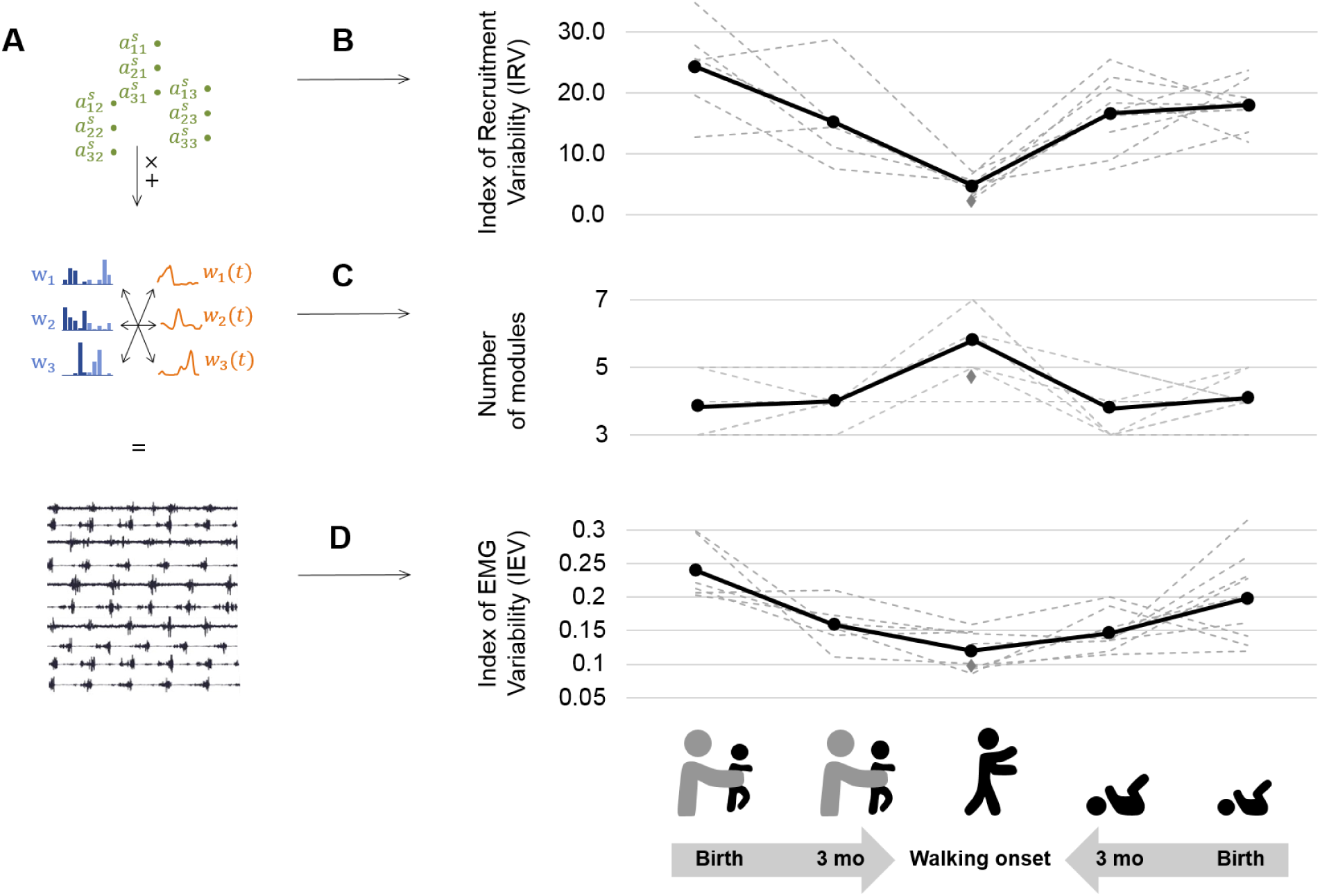
Decrease of variability between birth and walking onset associated with modifications of the underlying set of motor primitives. **A**. Computational elements contributing to the EMGs and their trial-to-trial variability (from top do down: activation coefficients, spatial and temporal modules, and muscle outputs). Graphs B C and D show how changes within the upper levels can explain the resulting motor variability during infant locomotor development. B. Variability of module activations, assessed by the index of recruitment variability (IRV). IRV represents the variability of the input that specifies which amplitude of activation has to be allocated to each possible pair of spatial and temporal modules. This index decreases from birth to walking onset considering stepping or kicking as neonatal behavior. C. Number of spatial and temporal modules, which increases from birth to walking onset considering stepping or kicking as neonatal behavior D. Index of EMG variability (IEV, same as in Figure 2I). This index decreases from birth to walking onset, considering stepping or kicking as neonatal behavior. The grey diamonds indicates the adult landmark.

### The number of motor primitives increases from birth to walking onset while variability decreases

A modular decomposition was applied to each dataset (for a given behavior, at a given age and for a given subject) thanks to NNMF. We found that several aspects of this decomposition was different depending on the age regarding both dimensionality (i.e. number of primitives) and variability of activations (Figure 3, Figure 4).

**Figure 4.**
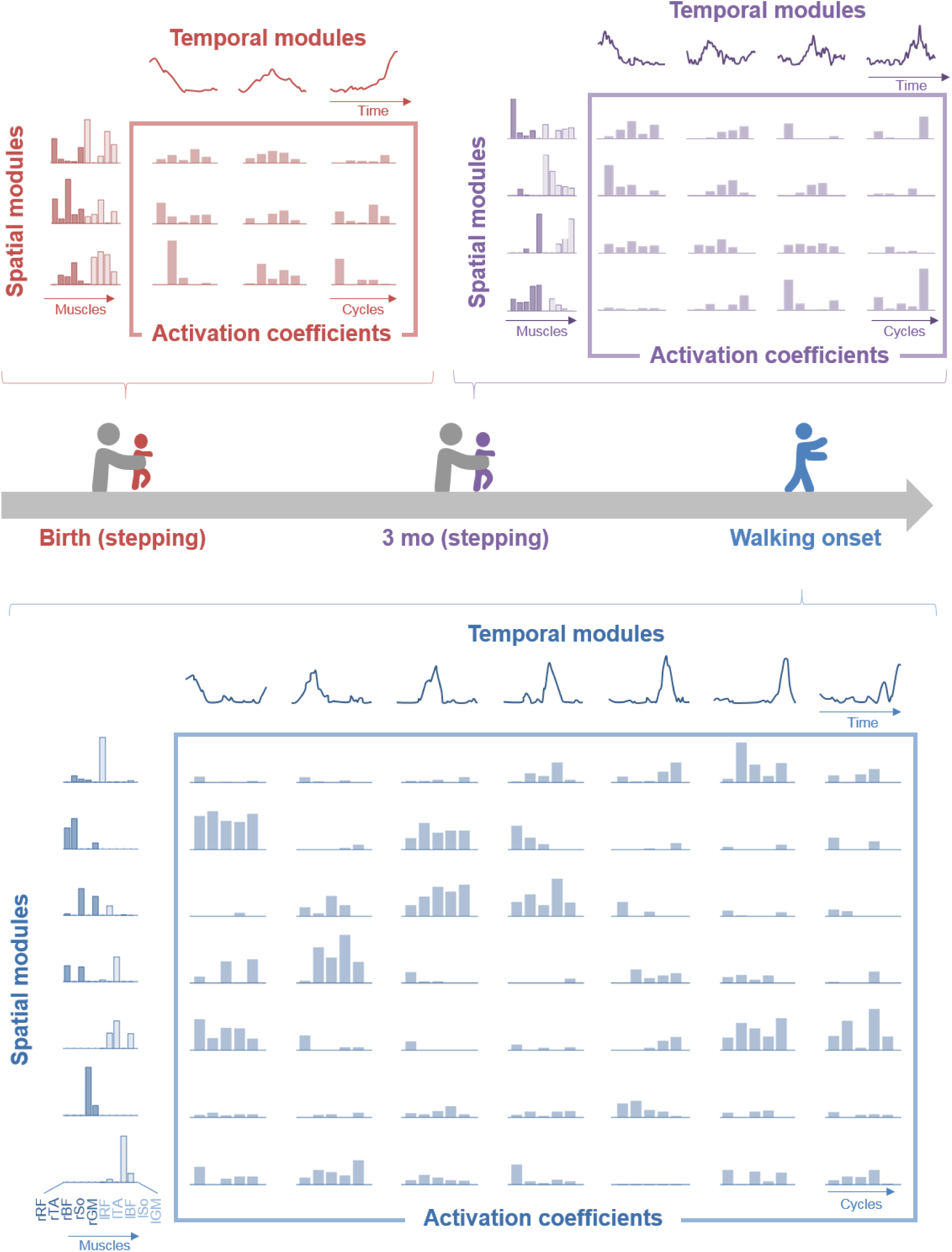
Modular organization at each age in a representative individual. At each age, EMG patterns can be decomposed into spatial modules and temporal modules (orange). Within each spatial module, weightings are plotted for muscles m1 to m10 in the following order: rectus femoris, tibialis anterior, biceps femoris, soleus, and gluteus medius (right muscles in dark colors followed by left muscles in light colors). Activation coefficients (at the crossing between each spatial and temporal modules) represent the level of activation of each possible pair of spatial and temporal modules during 5 steps. At birth (red, top left), EMG activity of stepping can be decomposed into 3 spatial and 3 temporal modules. At 3 mo (purple, top right), EMG activity of stepping can be decomposed on 4 spatial and 4 temporal modules. At walking onset (blue, bottom), EMG activity needs to be decomposed into 7 spatial and 7 temporal modules to get the same quality of modeling than at birth and at 3 mo with less modules. Activation coefficients are highly variable at birth and at 3 mo and less variable in toddlerhood, with some pairs that are nearly never activated across the 5 cycles. Note that toddler activations are still more variable than in adults (Figure 1).

To study dimensionality, we considered two approaches based on the Variance Accounted For (VAF) that is the index that indicates the quality of the modelling. The first approach identified the number of motor primitives (i.e. spatial and temporal modules) that are needed to reach a pre-determined VAF threshold. This threshold was established as the averaged VAF value identified in adult walking when extracting four spatial and temporal modules with the same algorithm, which is 0.75 (Hinnekens et al., 2020). This approach allowed to determine the number of modules of each individual and showed that the number of modules was more important at walking onset than at birth and at 3 mo (Figure 3C). The second approach set the number of modules to four as in standard adult walking, and relied on the analysis of the resulting VAF. This approach assessed dimensionality of the underlying modular system just like the first one but directly tested the hypothesis that four spatial and temporal modules are sufficient to adequately represent the given EMG signals across cycles. By relying on real numbers instead of integers, this second approach is useful because it is more suited to perform statistical analyses. It confirmed that a low-dimensional model fitted better at birth than at walking onset. We observed a significant VAF decrease between kicking at birth and walking (p=.047) and between kicking at 3 mo and walking (p=.016), as well as trends between stepping at birth and walking (p=.063) and between stepping at 3 mo and walking (p=.094). On average, the VAF for four spatial and temporal modules decreased with age, indicating that more and more modules were needed to equivalently reconstruct the EMG patterns, from birth and 3 mo to walking (Table S4). To sum up the modular organization was more complex in toddlers than in infants, as illustrated by Figure 4 and Figure 3B.

### Motor primitives are recruited with maximal variability during the neonatal period

After having analyzed the dimensionality of the signals, we wanted to explain how the IEV (EMG variability) could be higher in infants while their dimensionality was lower. Thus, we focused on the variability of activations of motor primitives. The Index of Recruitment Variability (IRV) which represents the extent to which spatial and temporal modules are variably activated across steps, significantly decreased in toddlers in comparison to infants (Figure 3A), indicating that module recruitment was less and less variable starting from either stepping or kicking from birth to walking (respectively p=.031 and p=.016) and from 3 mo to walking (respectively p=.031 and p=.016). To check that this effect was not due to differences in the number of modules we performed the same computations on values obtained by systematically extracting 4 spatial and temporal modules and found the same effects (Table S3). This shows that, even with the same number of modules, toddlers, almost like adults recruit modules in a more systematic way across cycles than infants (see Table S5 for individual data).

The Index of Recruitment Selectivity (IRS, which represents the extent to which a spatial module is always activated with the same temporal module and vice versa) remained quite stable across ages and behaviors, except between kicking at birth and walking when it showed a significant increase (p=.047). However, this index was always below the adult landmark at every age (Table S6), suggesting a low selectivity in the recruitment of spatial and temporal modules. Indeed a spatial module could be activated along with several temporal modules and vice versa depending on the cycle. We also repeated this computation after having extracted 4 spatial and temporal modules from each dataset and found the same results (see Table S5 for individual data).

### Motor primitives evolve between birth and walking onset toward gathering less muscles at a time

In order to identify if motor primitives would have been preserved across ages, we applied the best matching pairs method (d’Avella and Bizzi 2005) to our data and checked for similitudes between modules. As we could not find high similitudes we noticed that modules seemed to be less and less complex with time, suggesting a more individual muscle control (that is, spatial modules gathered the activation of less and less muscles, and temporal modules represented tighter and tighter peaks of activation, Figure 5A). To quantify this phenomenon, we created two indexes: the Selectivity of Muscular Activations Indexes (SMAI) and the Selectivity of Temporal Activation Indexes (STAI) (Table S5). The SMAI increased after 3 mo indicating that muscle weightings were sparser among spatial modules in toddlers than in infants. In other words, spatial modules were mostly composed of fewer muscles in toddlers compared to infants, (see Figure 5A and B). Interestingly, the value evolved in the opposite direction with respect to the adult landmark (Figure 5B). This increase occurred from stepping and kicking at birth to walking (respectively with a trend of p=.063 and significantly at p=.016) and significantly from stepping or kicking at 3 mo to walking (respectively p=.031 and p=.016). No effect was found between birth and 3 mo for stepping or kicking. Analyses were repeated on modular decompositions coming from the systematic extraction of 4 spatial and temporal modules and gave the same results (Table S3).

**Figure 5.**
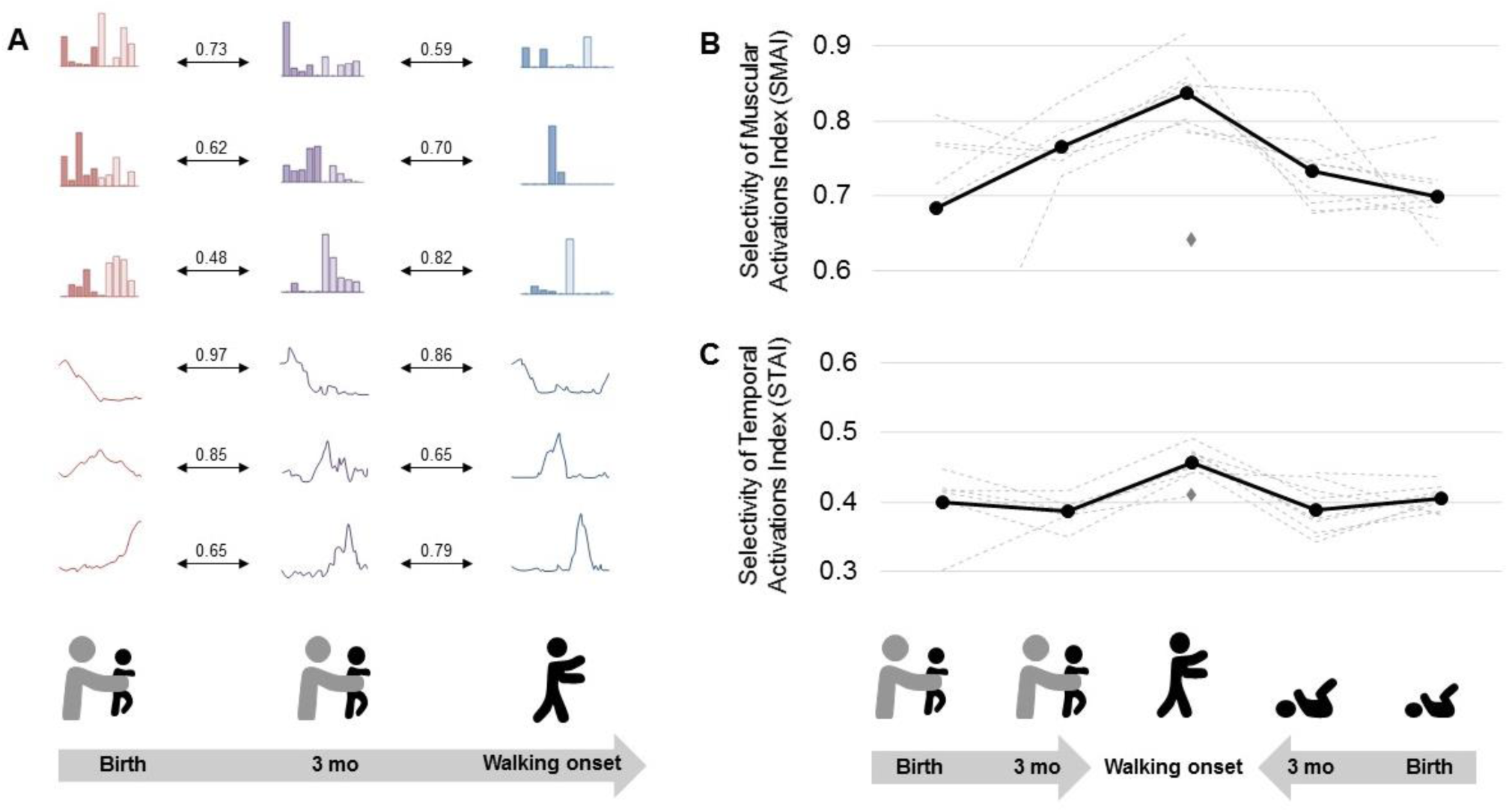
Development of modules’ structure from birth to walking onset. **A:** Similarity of modules between ages in a given individual according to the best matching pair method. **B:** Selectivity of muscular activations index (SMAI). **C:** Selectivity of temporal activations index (STAI). Both indexes increase between birth and walking onset considering stepping or kicking as neonatal behavior. The grey diamonds indicates the adult landmark.

The STAI also increased from birth and 3 mo to walking. This indicates that temporal modules had more tightened peaks of activation in toddlers than in infants (Figure 5A and C). The value for toddlers was also above the adult landmark (Figure 5C). For both stepping and kicking, the increase occurred between birth and walking (p=.09 and at p=.02 respectively) and between 3 mo and walking (respectively p=.03 and p=.02). Analyses were repeated following the extraction of 4 spatial and temporal modules and gave the same results (Table S3).

## Discussion

The objective of this study was to investigate whether and when trial-to-trial variability could emerge from the modular system of developing human infants. We found that the EMG variability of lower-limb muscles was maximal in the neonatal period and decreased at walking onset. This decrease of trial-to-trial variability was associated with an increase in the number of motor primitives. In the neonatal period the maximal EMG variability could be explained by variable activations and combinations of a small number of motor primitives, suggesting that the motor system can generate trial-to-trial variability at birth albeit relying on few primitives. In contrast at walking onset, more primitives were needed to explain EMG patterns of alternated leg movements, and these primitives were activated more consistently across trials. Furthermore, primitives were more selective than in infanthood, suggesting that toddlers developed the capacity to control less muscles at a time. Together, these results provide evidence that the motor system is flexible as soon as birth despite the existence of basic patterns of coordination, and suggest that neonatal motor primitives are plastic structures that fraction in early life to become more selective and more steadily activated. Below we discuss these findings in relation with the current literature about motor exploration and the maturation of the motor system through development.

This study highlights that EMG signals of alternated leg movements in human infants are highly variable as soon as birth, while neonatal behavior is often described as rather stereotyped due to the high prevalence of cocontractions (Dominici et al., 2011; Jeng et al., 2002; Spencer and Thelen, 2000; Thelen et al., 1981). Moreover, the signals analyzed here showed high variability while they were screened and selected among a lot of cycles that could be parallel or unilateral rather than alternate, which demonstrates even more flexibility than what we analyzed (Siekerman et al., 2015; Thelen et al., 1983, 1981). If we assume that trial-to-trial variability is a marker of motor exploration (Dhawale et al., 2017; Kao et al., 2008), our results would suggest that exploring the field of possible movements is possible as soon as birth despite the difficulty to uncouple the different muscles and joints of lower-limbs (Fetters et al., 2004; Teulier et al., 2012; Vaal et al., 2000). Such idea is coherent with the fact that motor learning abilities are in place very early in development and even before birth (Robinson, 2015; Robinson and Kleven, 2005).

Our results revealed that the most variable EMG output was seen at the time where the fewest number of primitives was found (i.e., in the neonatal period). This confirms and extends previous analyses relying on cross-sectional data showing that the motor system involves a few modules in rat and human newborns (Dominici et al., 2011). It also shows that variability in human can be generated through a structured modular system similarly to newborn rats (Blumberg et al., 2013), which seems to be specific to infanthood (as opposed to toddlerhood) according to our results. Although the neural interpretation is limited here, motor primitives were originally identified as spinal structures in animals (Bizzi et al., 1991) and are still hypothesized to be stored at a spinal level and activated through supraspinal inputs in adults (Bizzi and Cheung, 2013; Delis et al., 2014; Overduin et al., 2015, 2012; Roh et al., 2011). As such our results are reminiscent with neurophysiological studies in rodents that demonstrate that different motor patterns can emerge by applying different pharmacological or electrical stimuli to the neonatal spinal cord (Kiehn and Kjærulff, 1996; Klein et al., 2010) and that the neonatal motor system is able of flexibility via the variable activations of a few basic structures (Blumberg et al., 2013). Such variable activations could be due to the instability of the command and be incidental, but it could also be genuinely purposeful: in young birds learning to sing, the output appears to be highly variable, but becomes even more structured than the adult one after having inhibited some specific areas of the brain, proving that such areas were responsible for generating variability and facilitating exploration (Aronov et al., 2011). With growth, adaptive inhibitory mechanism settle in a way that does not compromise the capacity for future plasticity (Garst-Orozco et al., 2014). While similar neurophysiological investigations cannot be conducted in humans, our results shed new light about the potential existence of such mechanisms in human infants, as the motor output of neonates can also be decomposed into highly structured patterns and their temporary variable activations.

Strikingly, we observe the reverse phenomenon in toddlerhood: as infants grew to become toddlers, they seem to have widened their motor repertoire along with a stabilization of their command to produce alternate leg movements. Indeed more modules were needed to explain the EMG signals than in infanthood (Figure 3B), similarly to what was found in Dominici et al. (2011). The same trend was observed whether considering stepping or kicking, which were both shown to be neonatal locomotor precursors (Sylos-labini et al., 2020). The increase in the number of modules along the first year of life was associated with changes within the shape of modules themselves. They became more selective at walking onset (Figure 5), suggesting a more individual muscle control. This is in accordance with previous results reporting fractionation of primitives and decrease of duration of temporal modules (Sylos-labini et al., 2020) as well as consistent with the known decrease of cocontractions across the first year of life (Teulier et al., 2012). This is also strongly reminiscent with recent findings regarding the development of running, which starts with a fractionation of motor primitives between childhood and adulthood, allowing those primitives to be merged again latter with training (Cheung et al., 2020). On a different scale of analysis that was recently allowed by the use of high-density EMG in human neonates, it was also established that the synchronization across motoneurons of a given muscle was more important in neonates than in adults, which suggests a reshaping of central pattern generators during development (Del Vecchio et al., 2020). Assuming that modules are stored at a spinal level, such a reshaping within modules themselves is likely to be made early in infancy. Spinal circuits are indeed particularly plastic during early development thanks to the high activity of neurophysiological processes that can lead to changes within neurons excitability through practice (Brumley et al., 2015; Vinay et al., 2002). As neonates are able to set up several behaviors that involve flexion and extension cycles of lower-limbs, such as stepping and kicking but also air-stepping, crawling or swimming (Barbu-roth et al., 2014; Forma et al., 2019, 2018; McGraw, 1941, 1939), they could already benefit from a lot of opportunities to explore and to practice in order to shape the underlying circuits. The particular plasticity of motor primitives at this point is most likely allowed by the absence of stable synaptic connections within pathways, synaptogenesis being particularly active in the neonatal period (An et al., 2012; de Graaf-Peters and Hadders-algra, 2006).

The cross development between trial-to-trial variability of activations and the number of primitives suggests that the motor system is never designed to produce a maximal amount of variability. One could indeed picture a scenario where a maximal number of primitives would have been associated with maximally variable activations of these modules to maximize variability and exploration (or even no primitives at all to allow free muscle activations). However, constraining the space of possible options might be crucial for motor learning (Bernstein, 1967; Dhawale et al., 2017), which also relates to the exploration-exploitation tradeoff in reinforcement learning (Sutton and Barto, 1998). Humanoid robots indeed learn and build motor synergies more easily when starting to explore with a limited number of available degrees of freedom such as dynamic movement primitives (Lapeyre et al., 2011; Lungarella and Berthouze, 2003; Schaal, 2006). The developing system might have to always remain in an ideal compromise of constraint and flexibility, the difficulty of uncoupling lower-limb joints at the beginning of life (Fetters et al., 2004; Vaal et al., 2000) being an example of such helping constraint in humans. It was suggested that this rigid coupling that reduces the effective number of DOF allows to move with less need of processing capacities and without interferences of uncoordinated outputs (Piek, 2002). If this constraint might have a maturational origin, many factors could provide the same type of DOF reduction early in development (Newell et al., 1989), such as environmental ones (e.g. the space within the womb environment or the gravity at birth) and factors related to the task (e.g. kicking might allow to focus on lower-limbs only while walking also requires balance). Here we propose that the benefit of developmental motor primitives might be to yield an ideal space of possible that allow to efficiently explore several motor solutions via trial-to-trial variations of the activations of those temporary primitives. This might be critical to shape motor circuits, as suggested by the correlation between the lack of motor variability following early cerebral lesions and poor developmental outcomes in humans (Einspieler and Prechtl, 2005; Hadders-Algra, 2008). In this context, the approach proposed here could contribute to the quantitative study of motor variability in atypical development and characterize the extent to which motor exploration can be elicited in developing early rehabilitation protocols that are based on rhythmic behaviors (Angulo-Barroso et al., 2013; Campbell et al., 2012; Kolobe and Fagg, 2019; Sargent et al., 2020; Teulier et al., 2009).

When working with EMG to identify hypothetical motor primitives, several factors can limit interpretations such as EMG processing, cross-talk, or arbitrary choices made during EMG factorization. It is indeed worth noticing that the current study report different values than previous developmental studies (Dominici et al., 2011; Sylos-labini et al., 2020) regarding the absolute number of modules. However there is currently no consensus regarding the selection of the number of modules which depends on arbitrary criteria such as the VAF threshold. We can also note that Sylos-labini et al. (2020) reported more modules in kicking than in stepping at birth in contrast to what is reported here, however strict comparisons are limited on this point since we only studied supine kicking whereas these authors considered kicking both supine and vertically held. Also, to deal with the lack of consensus about EMG processing, we reproduced our analysis with other choices regarding filtering, amplitude normalization and time normalization, and verified that similar effects and trends emerged. This verification confirmed that our results were robust to reasonable processing choices. Cross-talk, which refers to the possibility of recording several muscles with one electrode, is particularly challenging in developmental studies since growth implies a modification of the distance across muscles. To deal with this issue we purposely chose muscles from different body regions (the 10 muscles are distributed from the hips to the shank and over the two lower-limbs). For agonist and antagonist muscles of a same body regions, we reproduced the analysis that was proposed by Dominici et al. (2011) ensuring that raw signals were not correlated (see Methods). Despite all precautions, the size of surface EMG electrodes remains a challenging issue of the field, and current development of high-density EMG might offer interesting perspectives for collecting more and more precise signals through surface EMG (Del Vecchio et al., 2020).

While we did not directly compare toddlers and adults data, adults landmarks suggested than walking in toddlers could rely on a higher dimensionality than adult walking (Figure 3B) involving more selective modules (i.e. modules controlling less muscles at a time, Figure 5B and 5C). Interestingly, it was recently suggested that several features of modularity could continue to evolve after toddlerhood: studying the development of walking and running in children, authors identified more primitives at 2 and 5 years old than in adults (Bach et al., 2021). As Cheung et al. (2020) showed that running motor primitives merge with training, we can hypothesize that walking primitives of toddlers will also merge with practice over time. Since learning to walk continues long after walking onset (Chang et al., 2006; Müller et al., 2013), it is not surprising to notice that the modular organization of toddlers could still need adjustments, even though it suggests that the system might first need to develop the capacity to control muscles more separately before gathering them again into more complex modules. As learning in a modular system relies on both learning the shape of modules and learning their activation parameters (d’Avella and Pai, 2010), the two processes might not be concomitant to ensure, as suggested above, to always remain in an ideal space of possibilities. Interestingly, recent data from rats report similar modular organization between organisms with different developmental history, suggesting that spinal primitives are determined early in development and conserved into adulthood (Yang et al., 2019). However, the fact that the neural repertoire is not mature before the important neural pruning of late adolescence in humans (de Graaf-Peters and Hadders-algra, 2006) suggests that module shaping could continue over a long period of time. The present study falls into this converging framework of long-lasting plasticity of motor modules, associating fractionation and merging of motor primitives throughout development and training (Bach et al., 2021; Cheung et al., 2020; Dominici et al., 2011; Hinnekens et al., 2020; Sylos-labini et al., 2020). While more studies are needed to characterize the maturation of the modular system after walking onset in humans, a long-lasting plasticity of motor primitives could allow to adapt to the development of the musculoskeletal system (Bizzi et al., 1991; Bizzi and Cheung, 2013) in order to integrate biomechanical specificities of each individual (Torres-Oviedo and Ting, 2010) as well as optimality considerations allowing the low energy costs of mature walking (Berret et al., 2019; Catavitello et al., 2018; de Rugy et al., 2012; Selinger et al., 2015).

## Materials and Methods

The protocol was in accordance with the Declaration of Helsinki and approved by the French Committee of People Protection. Families were recruited at the Port-Royal maternity in Paris. For each child, a parent provided informed written consent to participate to the study.

### Participants

12 infants (9 males 3 females) were tested longitudinally from birth to 3 months-old and 10 of them were tested shortly after they could walk independently. Human neonates with a few days of life being a rare clinical population, we choose the number of subject to replicate the one of Teulier et al. (2012) who analyzed the variability of EMG signal in a longitudinal follow-up of 12 infants between 1 months old and 12 months old in human infants. Inclusion criteria were no known physical or neurological disabilities, gestational age ≥ 38 weeks, weight ≥ 2800 g, and APGAR ≥ 8. The walking experiment was set up one to five weeks after infant would begin to use walking as the main mode of locomotion, based on parent’s reporting by phone call. They were also asked to write down the precise day when the child was able to “cross an entire room of about sixteen feet by walking”. One family forgot to write the precise date but still participated to the walking experiment. Two families participated until 3 months only and did not finish the study. Among the remaining toddlers, walking experience was 14.1 ± 7.9 days (mean ± SD) at the time of the experiment.

### Experimental Design

For each experiment, the first stage was to equip the infant or toddler with EMG sensors. Infants were then observed in several positions. At birth and at 3 months, they were observed in a supine position to observe kicking as well as held upright to induce stepping. We also observed two other behaviors that are not reported in this study (crawling and stepping on a treadmill). Each behavior was observed on a pediatric table during about 2 minutes in a randomized order for each infant and at each age. Regarding walking, toddlers were asked to go back and forth along a two meters exercise mat. They were walking barefoot at a natural speed during approximately one minute without any help from adults. Examples of each behavior are shown in Movie S1 (kicking at birth and around 3 months old in participant 3) and in Movie S2 (stepping at birth and around 3 months old in participant 4 followed by walking at walking onset).

### Data recording

#### EMG recording

Surface EMG data were recorded with the Cometa system at 2000 Hz. At all ages ten muscles from shanks thighs and buttocks were recorded bilaterally: the tibialis anterior, soleus, rectus femoris, biceps femoris and gluteus medius. Sensors were also placed on 6 other muscles of the trunk and the shoulders but the resulting data were not used in this study. The electrodes placement followed the SENIAM recommendations (Surface EMG for Non-Invasive Assessment of Muscles, seniam.org).

#### Video recording

We used 2D cameras (50Hz) at each age in order to detect cycles of flexion and extension, synchronized with the EMG recordings. Camera were placed on each side of the mat or the table so we would have a clear vision of both sides of the body to detect infants’ movements.

### Data processing and computed parameters

#### Identification of cycle events

Muscle modules are usually defined as invariant portions of the signal within step cycles. Here usual phases of stance and swing could have been defined for stepping and walking but not for kicking. As we wanted to be able to compare modules from different behaviors we choose to identify flexion and extension cycles instead of step cycles. Thus we identified two types of events in each behavior with two trained coders: beginning of hip flexion (BHF- defined as the first frame when hip flexion was elicited) and beginning of hip extension (BHE- defined as the first frame when hip extension was elicited). Reliability of the events identification by the coding procedure was excellent with an Intraclass Correlation Coefficient (ICC) of 0.99 (as done in Teulier et al., 2012).

A cycle was defined from a BHF to the following BHF, and made of two phases: the flexion phase, from BHF to BHE, and the extension phase, from BHE to BHF. We always analyzed cycles from the same side of the body, and kept this side for a given baby (for each behavior and each age). We only considered alternated cycles, defined as beginning between 10 and 90% of the cycle of the contralateral lower-limb. First and last cycles of an ensemble of alternating cycles were never considered. In order to compare basic kinematic parameters among behaviors we computed a few kinematic indexes from this coding of cycles: cycle duration, variability of step duration (standard deviation of step duration divided by the averaged step duration), and proportion of flexion and extension phases.

We retained data for our analysis only if a minimum of 5 alternated cycles of flexion and extension were made at both birth and at 3 months and were associated with a clean signal of the 10 muscles simultaneously. When more than 5 cycle were available we randomly selected the 5 cycles to include in the analysis. This conservative approach (i.e. selecting alternated cycles only) allowed to ensure reproducibility across ages and to focus only on the trial-to-trial variability that can be generated for a given behavior. Thus we got data for 9 infants for kicking, 6 infants for stepping and 10 toddlers for walking, with a total of 200 cycles included in the analysis (Table S1).

#### EMG processing

EMG data are presented on Figure 2 and the procedure for EMG processing is presented on Figure S1. For each signal we applied a high pass filter (40hz, fourth order Butterworth filter) followed by a rectification, as in Ivanenko (2013). To smooth envelopes we used a moving median since some artefacts were often visible in kicking due to the leg touching the other leg or the mat. The window of this moving median was normalized relatively to the cycle duration that changed with the age of the Infant, as described in Ivanenko (2013). As cycle duration was about twice lower on average in 3-month old infants, and again twice lower in toddlers, the moving median window was 400 points in newborns, 200 points in 3- month olds and 100 points in toddlers. This filtering procedure was applied to the entire recordings of stepping, kicking and walking of each baby in order to proceed with amplitude normalization. Regarding toddler, the amplitude normalization was made under the maximal value of the whole walking recording. Finally, cycles were isolated and normalized in time so that each cycle would correspond to 200 time points, by interpolating the flexion phase to 80 points and the extension phase to 120 points, based on the proportion of phases of independent walking. The rationale here is that all cycles will temporally match the same kinematic events regardless of the age or the behavior. To test how choices on amplitude and time normalization could affect our results, we repeated the analyses of this paper with a different amplitude normalization (normalizing each ensemble of 5 cycles under its own maximum) and with a different time normalization (interpolating to the cycle without fixing the phases), which confirmed that our results were not dependent of those methodological choices.

As cross-talk might be an issue when recording surface EMG data, we used the same criterion than Dominici et al. (2011) to assess potential cross-talk (Pearson correlation coefficient >0.2 among pairs of agonist and antagonist muscles). At birth, 3% of our sample had a correlation coefficient >0.2; 1.4% at 3 months, and 6% at walking onset. For these samples, we checked whole recordings and verified that different strides from one subject were not all >0.2.

#### Variability of EMG signals

In order to compare the variability of EMG signals across ages, we computed an Index of EMG Variability (Hinnekens et al., 2020) from these processed EMG signals, as the standard deviation computed point by point across the 5 cycles. This allowed to characterize the motor variability associated with each behavior and at each age before trying to explain how this variability could be generated within a modular system.

#### EMG factorization

We extracted spatial and temporal muscle modules into the EMG signals thanks to the Space-by-Time Decomposition method. This method uses non-negative matrix factorization (NNMF) in order to factor EMG signal in a given number of invariant components. As described in Delis et al. (2014) this method unifies previous ones as it allows to identify both spatial and temporal EMG modules. In addition, it allows to preserve intra-individual variability into the analysis, through activation coefficients that represent the degree of activation of a given module in each cycle. An EMG signal is considered as a double linear combination of invariant spatial and temporal modules, so that any muscle pattern ***m***_*s*_*(t)* of the cycle scan be written:

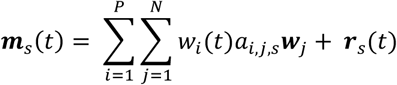

where *P* and *N* are the numbers of temporal and spatial modules respectively, *w*_*i*_*(t)* and ***w***_*j*_ are the temporal and spatial modules respectively, *a*_*i,j,s*_ is a scalar activation coefficient (function of the pair of modules it activates and step s), and ***r***_*s*_*(t)* is the residual reconstruction error describing the difference between the original signal and the reconstructed one. A spatial module is a 10-dimensional vector (as we recorded 10 muscles) and describes ensemble group of muscles that are invariantly activated together across cycles with the same relative proportions. A temporal module is a time-varying function (here described by a 200- dimensional vector) and represents the invariant activation timing of a spatial module within a cycle. The relative shape of spatial and temporal module are both considered invariant across cycles, but the way pairs of spatial and temporal modules are activated together can differ from one cycle to another. To represent those potentially variable activations, the method computes scalar activation coefficients for each cycle and for each possible pair of spatial and temporal modules, thereby implementing a dimensionality reduction in EMG space. A high activation coefficient corresponds to the concurrent activation of z a specific pair of spatial and temporal modules. The algorithm finds the best fit of spatial and temporal modules and activation coefficients by progressively modifying their values until reaching a convergence criterion. The extraction procedure is repeated 50 times to prevent the risk of finding non-optimal values because of local minima. This algorithm is detailed in Delis et al. (2014). It was run with a custom Matlab® code. While the goal of this procedure is to model the underlying motor command, we cannot presume about the neural origin of the identified modules, thus we sometimes refer to the identified modular organization as modularity or dimensionality “within the motor output” in the method and result sections.

#### Goodness of fit criteria

We computed the VAF (Variance Accounted For) as a quality of reconstruction criteria. The VAF is the coefficient of determination between the initial matrix and the reconstructed:

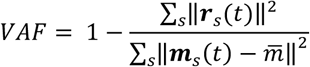

Where 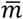 is the mean level of muscle activity across all samples and ‖ ‖ represents the Frobenius norm.

The VAF quantifies the goodness of fit between the original EMG patterns and those that are reconstructed from the decomposition. The usual approach is to extract different numbers of modules from each initial matrix and to choose the number that allows to get a preset threshold VAF. Nonetheless it is also interesting to compare the VAF for a given number of module as it allows to directly test the extent to which a low-dimensional command model can give a faithful description of the initial EMG patterns. Moreover the first approach can only give integers while the second assesses the dimensionality of the motor output by returning reals which might be more precise. Thus we used here both approaches due to their complementarity.

For the first approach, which is to establish a module number given a threshold VAF, we used the threshold of 0.75. The rationale is that numerous studies showed that adults walking could be efficiently modelled by four spatial and temporal modules that are biomechanically functional (Clark et al., 2010; Lacquaniti et al., 2012; Neptune et al., 2009) and that the average VAF of 0.75 was obtained when using the space-by-time decomposition method on non-averaged signals of adults (Hinnekens et al., 2020). Thus we considered this value to represent an efficient modularity with this methodology and used it as a critical threshold. We repeated the extraction and increased the number of spatial and temporal modules until crossing this threshold. For simplicity, we only considered scenarios where the number of spatial modules and the number of temporal modules were equivalent (P=N). This approach allowed us to determine a necessary number of modules in each task and each age.

For the second approach which is to establish the VAF for a given number of modules, we again relied on previous results on the EMG decomposition in adults and computed the VAF corresponding to four spatial and temporal modules for each task and each age.

Following those computations we used the number of modules identified with the first approach to compute the metrics presented below. Nevertheless those metrics could be influenced by differences in module numbers, thus they were also all computed for 4 spatial and 4 temporal modules as a control setting to ensure that significant differences among those metrics would not just be due to differences in the dimensionality of each decomposition.

#### Indexes describing the recruitment of modules]

We computed two indexes to describe how the recruitment of modules would evolve across ages: the Index of Recruitment Variability (IRV) and the Index of Recruitment Selectivity (IRS).

The IRV is computed as the average standard deviation of activation coefficients across cycles. As such it quantifies the variability of modules recruitment: a high value means that modules are differently recruited across cycles, while a low value indicates a stable recruitment of modules across cycles.

The IRS corresponds to the sparseness of activation coefficients, using a metric described in Hoyer (2004). It quantifies the selectivity of activations between spatial and temporal modules: the larger IRS is, the more spatial and temporal modules are exclusively paired, while the lower IRS is, the more spatial and temporal modules are multiplexed.

#### Indexes describing the nature of modules themselves

We computed two additional indexes to describe how the nature of modules would evolve across ages: the Selectivity of Muscular Activations Index (SMAI) and the Selectivity of Temporal Activations Index (STAI).

The SMAI is computed as the average sparseness of spatial modules. Spatial modules are 10- dimensional weighting vectors that can be composed mainly of one muscle (single-muscle module) or ten muscles with equivalent weights (co-activation module). In general, within a spatial module, muscle will be weighted with more or less. The SMAI quantifies how muscle-selective is a spatial module: when the SMAI is large, it means that spatial modules only gather a few muscles (and in the extreme case, the spatial module is formed with only one muscle) while when it is low, it means that spatial modules gather many different muscles with non-negligible weights.

The STAI is computed as the average sparseness of the temporal activations across temporal modules. The peaks of activations within temporal modules could indeed be narrow (high STAI values), describing refined and precise temporal activation across a cycle, or spread (low STAI values), describing a more continuous activation across the large part of the cycle.

#### Computing adult values for comparisons

When analyzing the results it is often useful to know the value that would be obtained for adults regarding our parameters of interest. Thus we used data from 12 adults from another study (Hinnekens et al., 2020) to compute similar indexes on 5 step cycles per adult. No statistic were made from these data but values were depicted on figures to show the adult landmark (Figure 1, grey diamonds on Figure 4 and Figure 5, Table S6).

### Statistical analyses

In addition to reporting the goodness of fit associated with our modeling (i.e. VAF value), we compared features of variability and modularity across ages. Because the number of participants was small we used non-parametric Wilcoxon tests to compare paired samples across ages from each of the two precursor behaviors to walking on basic kinematic parameters (cycle duration, variability of cycle duration and proportion of flexion and extension phases across a cycle) and on the variability of EMG signals (IEV). With the same test we analyzed the dimensionality of the motor output by comparing the VAF for a fixed number of modules as well as the number of necessary modules to reach a given VAF. Finally, we compared indexes describing the recruitment of modules (IRV and IRS) as well as indexes describing the nature of modules themselves (SMAI and STAI). Statistical p-values are summarized in Table S3.

## Acknowledgments

We thank Prof. François Goffinet, head of the Port-Royal maternity in Paris, for encouraging this study. We also thank the Région Ile-de-France for their participation in the initial set-up of the Babylab. We thank Dr Sophie Bavard for insightful comments on the manuscript. Finally we warmly thank all infants and parents who participated in the study.

## References

An S, Yang J, Sun H, Kilb W, Luhmann HJ. 2012. Long-Term Potentiation in the Neonatal Rat Barrel Cortex In Vivo. J Neurosci 32:9511–9516. doi:10.1523/JNEUROSCI.1212-12.2012.

Angulo-Barroso RM, Tiernan C, Chen LC, Valentin-Gudiol M, Ulrich DA. 2013. Treadmill training in moderate risk preterm infants promotes stepping quality-Results of a small randomised controlled trial. Res Dev Disabil 34:3629–3638. doi:10.1016/j.ridd.2013.07.037

Aronov D, Veit L, Goldberg JH, Fee MS. 2011. Two distinct modes of forebrain circuit dynamics underlie temporal patterning in the vocalizations of young songbirds. J Neurosci 31:16353–16368. doi:10.1523/JNEUROSCI.3009-11.2011

Bach MM, Daffertshofer A, Dominici N. 2021. Muscle Synergies in Children Walking and Running on a Treadmill. Front Hum Neurosci 15. doi:10.3389/fnhum.2021.637157

Barbu-roth M, Anderson DI, Despre A, Streeter RJ, Cabrol D, Trujillo M, Campos JJ. 2014. Air Stepping in Response to Optic Flows That Move Toward and Away From the Neonate. Dev Psychobiol 56:1142–1149. doi:10.1002/dev.21174

Bernstein NA. 1967. The co-ordination and regulation of movements, 1st ed. Oxford.

Berret B, Delis I, Gaveau J, Jean F. 2019. Optimality and Modularity in Human Movement: From Optimal Control to Muscle Synergies In: Venture G, Laumond J-P, Watier B, editors. Biomechanics of Anthropomorphic Systems. Springer, Cham. pp. 105–133. doi:10.1007/978-3-319-93870-7

Bizzi E, Cheung VCK. 2013. The neural origin of muscle synergies. Front Comput Neurosci 7:1–6. doi:10.3389/fncom.2013.00051

Bizzi E, Cheung VCK, d’Avella A, Saltiel P, Tresch M. 2008. Combining modules for movement. Brain Res Rev 57:125–133. doi:10.1016/j.brainresrev.2007.08.004

Bizzi E, Mussa-Ivaldi F, Giszter S. 1991. Computations underlying the execution of movement: a biological perspective. Science 253:287–291. doi:10.1126/science.1857964

Blumberg MS, Coleman CM, Gerth AI, McMurray B. 2013. Spatiotemporal structure of REM sleep twitching reveals developmental origins of motor synergies. Curr Biol 23:2100–2109. doi:10.1016/j.cub.2013.08.055

Brumley MR, Kauer SD, Swann HE. 2015. Developmental plasticity of coordinated action patterns in the perinatal rat. Dev Psychobiol 57:409–420. doi:10.1002/dev.21280

Campbell SK, Gaebler-Spira D, Zawacki L, Clark A, Boynewicz K, Deregnier RA, Kuroda MM, Bhat R, Yu J, Campise-Luther R, Kale D, Bulanda M, Zhou XJ. 2012. Effects on motor development of kicking and stepping exercise in preterm infants with periventricular brain injury: A pilot study. J Pediatr Rehabil Med 5:15–27. doi:10.3233/PRM-2011-0185

Catavitello G, Ivanenko Y, Lacquaniti F. 2018. A kinematic synergy for terrestrial locomotion shared by mammals and birds. Elife 7:1–28. doi:10.7554/eLife.38190

Chang C-L, Kubo M, Buzzi U, Ulrich BD. 2006. Early changes in muscle activation patterns of toddlers during walking. Infant Behav Dev 29:175–88. doi:10.1016/j.infbeh.2005.10.001

Cheung VCK, Cheung BMF, Zhang JH, Chan ZYS, Ha SCW, Chen CY, Cheung RTH. 2020. Plasticity of muscle synergies through fractionation and merging during development and training of human runners. Nat Commun 11:1–15. doi:10.1038/s41467-020-18210-4

Clark DJ, Ting LH, Zajac FE, Neptune RR, Kautz SA. 2010. Merging of Healthy Motor Modules Predicts Reduced Locomotor Performance and Muscle Coordination Complexity Post-Stroke. J Neurophysiol 103:844–857. doi:10.1152/jn.00825.2009

Cohn BA, Szedlák M, Gärtner B, Valero-Cuevas FJ. 2018. Feasibility Theory Reconciles and Informs Alternative Approaches to Neuromuscular Control. Front Comput Neurosci 12:1–18. doi:10.3389/fncom.2018.00062

d’Avella A, Pai DK. 2010. Modularity for sensorimotor control: Evidence and a new prediction. J Mot Behav 42:361–369. doi:10.1080/00222895.2010.526453

d’Avella A, Saltiel P, Bizzi E. 2003. Combinations of muscle synergies in the construction of a natural motor behavior. Nat Neurosci 6:300–308. doi:10.1038/nn1010

de Graaf-Peters VB De, Hadders-algra M. 2006. Ontogeny of the human central nervous system : What is happening when ? Early Hum Dev 82:257–266. doi:10.1016/j.earlhumdev.2005.10.013

de Rugy A, Loeb GE, Carroll TJ. 2012. Muscle Coordination Is Habitual Rather than Optimal. J Neurosci 32:7384–7391. doi:10.1523/JNEUROSCI.5792-11.2012

Del Vecchio A, Sylos-Labini F, Mondì V, Paolillo P, Ivanenko Y, Lacquaniti F, Farina D. 2020. Spinal motoneurons of the human newborn are highly synchronized during leg movements. Sci Adv.

Delis I, Panzeri S, Pozzo T, Berret B. 2014. A unifying model of concurrent spatial and temporal modularity in muscle activity. J Neurophysiol 111:675–93. doi:10.1152/jn.00245.2013

Dhawale AK, Smith MA, Ölveczky BP. 2017. The Role of Variability in Motor Learning. Annu Rev Neurosci 40:479–498. doi:10.1146/annurev-neuro-072116-031548

Dominici N, Ivanenko YP, Cappellini G, D’Avella A, Mondì V, Cicchese M, Fabiano A, Silei T, Di Paolo A, Giannini C, Poppele RE, Lacquaniti F. 2011. Locomotor primitives in newborn babies and their development. Science 334:997–9. doi:10.1126/science.1210617

Einspieler C, Prechtl HFR. 2005. Prechtl’s Assessment of General Movements: A Diagnostic Tool for the Functional Assessment of the Young Nervous System. Ment Retard Dev Disabil Res Rev 11:61–67. doi:10.1002/mrdd.20051

Fetters L, Chen YP, Jonsdottir J, Tronick EZ. 2004. Kicking coordination captures differences between full-term and premature infants with white matter disorder. Hum Mov Sci 22:729–748. doi:10.1016/j.humov.2004.02.001

Forma V, Anderson DI, Goffinet F, Barbu-Roth M. 2018. Effect of optic flows on newborn crawling. Dev Psychobiol 60:497–510. doi:10.1002/dev.21634

Forma V, Anderson DI, Huet V, Granjon L, Barbu-roth M. 2019. What Does Prone Skateboarding in the Newborn Tell Us About the Ontogeny of Human Locomotion ? Child Dev 90:1286–1302. doi:10.1111/cdev.13251

Garst-Orozco J, Babadi B, Ölveczky BP. 2014. A neural circuit mechanism for regulating vocal variability during song learning in zebra finches. Elife 3:e03697. doi:10.7554/eLife.03697

Hadders-algra M. 2018. Early human motor development: From variation to the ability to vary and adapt. Neurosci Biobehav Rev 90:411–427. doi:10.1016/j.neubiorev.2018.05.009

Hadders-Algra M. 2008. Reduced variability in motor behaviour: An indicator of impaired cerebral connectivity? Early Hum Dev 84:787–789. doi:10.1016/j.earlhumdev.2008.09.002

Hinnekens E, Berret B, Do M, Teulier C. 2020. Modularity underlying the performance of unusual locomotor tasks inspired by developmental milestones. J Neurophysiol 123:496–510.

Hoyer PO. 2004. Non-negative Matrix Factorization with Sparseness Constraints. J Mach Learn Res 5:1457–1469.

Ivanenko YP, Poppele RE, Lacquaniti F. 2004. Five basic muscle activation patterns account for muscle activity during human locomotion. J Physiol 556:267–82. doi:10.1113/jphysiol.2003.057174

Jeng SF, Chen LC, Yau KIT. 2002. Kinematic analysis of kicking movements in preterm infants with very low birth weight and full-term infants. Phys Ther 82:148–159. doi:10.1093/ptj/82.2.148

Kao MH, Wright BD, Doupe AJ. 2008. Neurons in a forebrain nucleus required for vocal plasticity rapidly switch between precise firing and variable bursting depending on social context. J Neurosci 28:13232–13247. doi:10.1523/JNEUROSCI.2250-08.2008

Kiehn O, Kjærulff O. 1996. Spatiotemporal characteristics of 5-HT and dopamine-induced rhythmic hindlimb activity in the in vitro neonatal rat. J Neurophysiol 75:1472–1482. doi:10.1152/jn.1996.75.4.1472

Klein DA, Patino A, Tresch MC. 2010. Flexibility of Motor Pattern Generation Across Stimulation Conditions by the Neonatal Rat Spinal Cord. J Neurophysiol 103:1580–1590. doi:10.1152/jn.00961.2009.

Kolobe THA, Fagg AH. 2019. Robot Reinforcement and Error-Based Movement Learning in Infants with and Without Cerebral Palsy. Phys Ther 99:677–688. doi:10.1093/ptj/pzz043

Lacquaniti F, Ivanenko YP, Zago M. 2012. Patterned control of human locomotion. J Physiol 590:2189–2199. doi:10.1113/jphysiol.2011.215137

Lapeyre M, Ly O, Oudeyer P, Lapeyre M, Ly O, Maturational PO, Lapeyre M, Ly O. 2011. Maturational constraints for motor learning in high-dimensions : the case of biped walking 2011 11th IEEE-RAS International Conference on Humanoid Robots. pp. 707–714.

Lungarella M, Berthouze L. 2003. On the interplay between morphological, neural, and environmental dynamics: A robotic case study. Adapt Behav 10:223–241. doi:10.1177/1059712302010003005

Mandelblat-Cerf Y, Paz R, Vaadia E. 2009. Trial-to-trial variability of single cells in motor cortices is dynamically modified during visuomotor adaptation. J Neurosci 29:15053–15062. doi:10.1523/JNEUROSCI.3011-09.2009

McGraw MB. 1941. Development of neuro-muscular mechanisms as reflected in the crawling and creeping behavior of the human infant. J Genet Psychol 83–111.

McGraw MB. 1939. Swimming behavior of the human infant. J Pediatr 15:485–490. doi:10.1016/S0022-3476(39)80003-8

Müller J, Müller S, Baur H, Mayer F. 2013. Intra-individual gait speed variability in healthy children aged 1-15 years. Gait Posture 38:631–636. doi:10.1016/j.gaitpost.2013.02.011

Neptune RR, Clark DJ, Kautz SA. 2009. Modular control of human walking : A simulation study. J Biomech 42:1282–1287. doi:10.1016/j.jbiomech.2009.03.009

Newell KM, Van Emmereik REA, McDonald PV. 1989. Biomechanical Constraints and Action Theory. Hum Mov Sci 8:403–409.

Overduin SA, d’Avella A, Carmena JM, Bizzi E. 2012. Microstimulation Activates a Handful of Muscle Synergies. Neuron 76:1071–1077. doi:10.1016/j.neuron.2012.10.018

Overduin SA, d’Avella A, Roh J, Carmena JM, Bizzi E. 2015. Representation of muscle synergies in the primate brain. J Neurosci 35:12615–12624. doi:10.1523/JNEUROSCI.4302-14.2015

Piek JP. 2002. The role of variability in early motor development. Infant Behav Dev 25:452–465. doi:10.1016/S0163-6383(02)00145-5

Robinson SR. 2015. Spinal mediation of motor learning and memory in the rat fetus. Dev Psychobiol 57:421–434. doi:10.1002/dev.21277

Robinson SR, Kleven GA. 2005. Learning to move before birth. Prenat Dev postnatal Funct.

Roh J, Cheung VCK, Bizzi E. 2011. Modules in the brain stem and spinal cord underlying motor behaviors. J Neurophysiol 106:1363–1378. doi:10.1152/jn.00842.2010

Sargent B, Havens K, Wisnowki J, Wu T-W, Kubo M, Fetters L. 2020. In-home kicking-activated mobile task to motivate selective motor control of infants at high risk of cerebral palsy: a feasibility study. Phys Ther 100:2217–2226.

Schaal S. 2006. Dynamic Movement Primitives -A Framework for Motor Control in Humans and Humanoid Robotics. Adapt Motion Anim Mach 261–280. doi:10.1007/4-431-31381-8_23

Selinger JC, O’Connor SM, Wong JD, Donelan JM. 2015. Humans Can Continuously Optimize Energetic Cost during Walking. Curr Biol 25:2452–2456. doi:10.1016/j.cub.2015.08.016

Siekerman K, Barbu-Roth M, Anderson DI, Donnelly A, Goffinet F, Teulier C. 2015. Treadmill stimulation improves newborn stepping. Dev Psychobiol 57:247–254. doi:10.1002/dev.21270

Spencer JP, Thelen E. 2000. Spatially Specific Changes in Infants’ Muscle Coactivity as They Learn to Reach. Infancy 1:275–302. doi:10.1207/S15327078IN0103_1

Sutton RS, Barto AG. 1998. Reinforcement Learning, MA: MIT Pr. ed. Cambridge.

Sylos-labini F, Scaleia V La, Cappellini G, Fabiano A, Picone S, Keshishian ES, Zhvansky DS, Paolillo P, Irina A, D’Avella A, Ivanenko YP, Lacquaniti F, Solopova IA, D’Avella A, Ivanenko YP, Lacquaniti F. 2020. Distinct locomotor precursors in newborn babies. Proc Natl Acad Sci 117:9604–9612. doi:10.1073/pnas.1920984117

Teulier C, Sansom JK, Muraszko K, Ulrich BD. 2012. Longitudinal changes in muscle activity during infants’ treadmill stepping. J Neurophysiol 108:853–862. doi:10.1152/jn.01037.2011

Teulier C, Smith BA, Kubo M, Chang C-L, Moerchen V, Murazko K, Ulrich BD. 2009. Stepping Responses of Infants With Myelomeningocele When Supported on a Motorized Treadmill. Phys Ther 89:60–72. doi:10.2522/ptj.20080120

Thelen E, Bradshaw G, Ward JA. 1981. Spontaneous kicking in month-old infants: Manifestation of a human central locomotor program. Behav Neural Biol 32:45–53. doi:10.1016/S0163-1047(81)90257-0

Thelen E, Ridley-johnson R, Fisher DM. 1983. Shifting Patterns of Bilateral Coordination and Lateral Dominance in the Leg Movements of Young Infants. Dev Psychobiol 16:29–46.

Torres-Oviedo G, Ting LH. 2010. Subject-Specific Muscle Synergies in Human Balance Control Are Consistent Across Different Biomechanical Contexts. J Neurophysiol 103:3084–3098. doi:10.1152/jn.00960.2009

Tresch MC, Saltiel P, Bizzi E. 1999. The construction of movement by the spinal cord. Nat Neurosci 2:162–167. doi:10.1038/5721

Vaal J, Van Soest AJ, Hopkins B, Sie LTL, Van der Knaap MS. 2000. Development of spontaneous leg movements in infants with and without periventricular leukomalacia. Exp Brain Res 135:94–105. doi:10.1007/s002210000508

Valero-Cuevas FJ. 2009. A Mathematical Approach to the Mechanical Capabilities of Limbs and Fingers In: Sternad D, editor. Progress in Motor Control. Springer, Boston, MA. pp. 619–633. doi:10.1007/978-0-387-77064-2

Vinay L, Brocard F, Clarac F, Norreel JC, Pearlstein E, Pflieger JF. 2002. Development of posture and locomotion: An interplay of endogenously generated activities and neurotrophic actions by descending pathways. Brain Res Rev 40:118–129. doi:10.1016/S0165-0173(02)00195-9

Yang Q, Logan D, Giszter SF. 2019. Motor primitives are determined in early development and are then robustly conserved into adulthood. Proc Natl Acad Sci U S A 116. doi:10.1073/pnas.1821455116

