## Supplemental Figure 1 for "Generating variability from motor primitives during infant locomotor development"

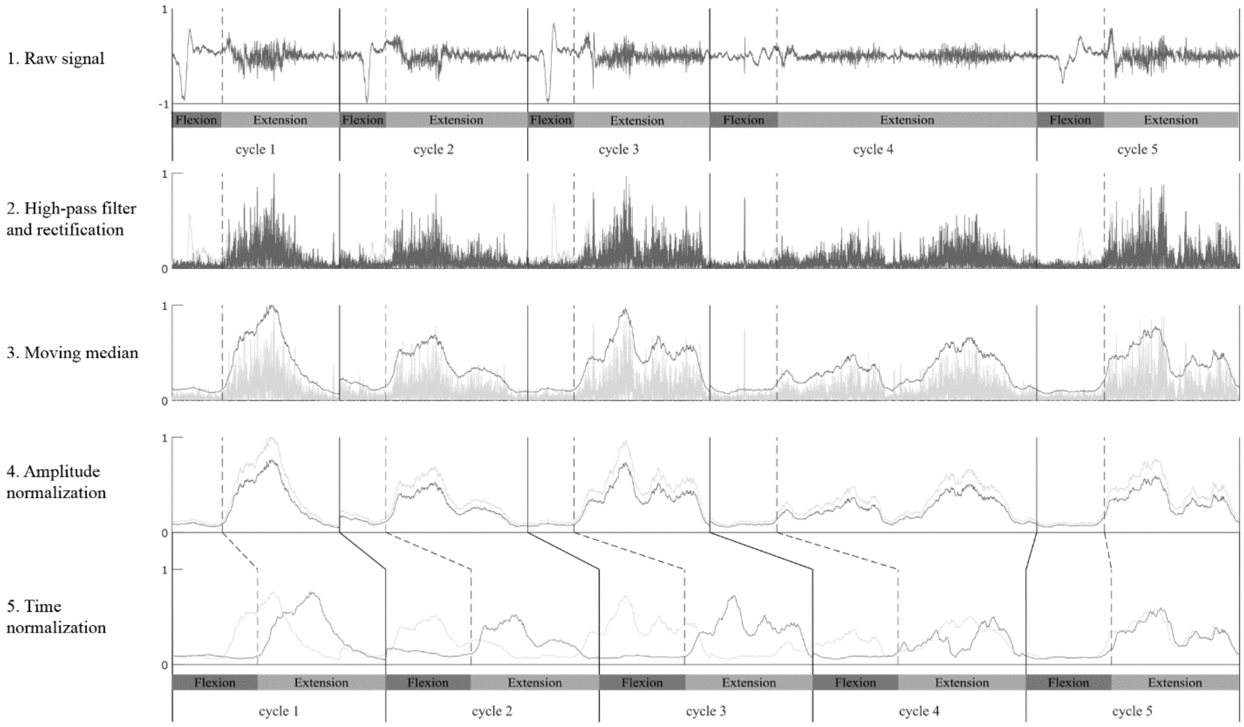


**Figure S1.**

Processing of EMG data. Each line shows one step of the preprocessing for an ensemble of 5 flexion and extension cycles for one muscle. On each graph, the dark signal represent the named step and the lighter signal the previous step (i.e. from the above line).
