## Supplemental Table 1 for "Generating variability from motor primitives during infant locomotor development"

**Table S1.**
Summary of individual characteristics.

| Subject ID | Gender | Birth Weight (kg) | Birth High (cm) | Recorded precursor | Age in days | | |
|---|---|---|---|---|---|---|---|
| | | | | | Around birth | Around 3 months | Around walking onset |
| 1 | F | 3.71 | 52 | Stepping | 3 | 87 | 412 |
| 2 | M | 3.46 | 48 | Kicking | 2 | 98 | 371 |
| 3 | F | 3.07 | 50 | Kicking | 2 | 86 | / |
| 4 | M | 3.74 | 51 | Stepping & Kicking | 2 | 74 | 434 |
| 5 | M | 2.99 | 50 | Stepping & Kicking | 2 | 79 | 365 |
| 6 | M | 3.47 | 51 | Kicking | 1 | 90 | 409 |
| 7 | M | 3.39 | 51 | Kicking | 2 | 122 | 478 |
| 8 | M | 3.66 | 49 | Stepping & Kicking | 2 | 117 | 536 |
| 9 | M | 3.28 | 48.5 | Stepping | 21 | 82 | 446 |
| 10 | F | 3.89 | 51 | Stepping | 2 | 116 | 428 |
| 11 | M | 3.55 | 52 | Kicking | 2 | 120 | 380 |
| 12 | M | 3.76 | 51 | Kicking | 8 | 105 | / |
