## Supplemental Table 2 for "Generating variability from motor primitives during infant locomotor development"

**Table S2.**

Individual data regarding basic EMG and kinematic parameters (Figures 2F to 2H and 4D).

| Precursor | Subject ID | Cycle duration (s) | | | Proportion of extension phase (%) | | |
|---|---|---|---|---|---|---|---|
| | | Birth | 3 months | Walking | Birth | 3 months | Walking |
| **Stepping** | 1 | 3.56 | 3.37 | 0.70 | 56.34 | 61.90 | 72.58 |
| | 4 | 5.07 | 1.95 | 0.97 | 74.59 | 73.11 | 72.55 |
| | 5 | 2.77 | 1.41 | 0.66 | 73.21 | 70.94 | 71.79 |
| | 8 | 2.79 | 1.93 | 0.66 | 45.71 | 54.78 | 71.00 |
| | 9 | 4.74 | 2.63 | 0.66 | 80.92 | 64.61 | 68.89 |
| | 10 | 5.39 | 2.92 | 0.94 | 70.67 | 78.93 | 76.55 |
| | **Mean** | **4.05** | **2.37** | **0.77** | **66.91** | **67.38** | **72.23** |
| | **SD** | **1.16** | **0.73** | **0.15** | **13.19** | **8.66** | **2.52** |
| **Kicking** | 2 | 3.43 | 0.52 | 0.73 | 68.28 | 44.04 | 70.47 |
| | 3 | 3.77 | 0.65 | / | 44.92 | 56.89 | / |
| | 4 | 2.63 | 1.59 | 0.97 | 63.33 | 48.21 | 72.55 |
| | 5 | 4.24 | 1.66 | 0.66 | 66.81 | 55.78 | 71.79 |
| | 6 | 1.68 | 1.42 | 0.96 | 55.90 | 49.92 | 79.69 |
| | 7 | 5.86 | 0.60 | 0.94 | 80.65 | 43.55 | 74.26 |
| | 8 | 1.97 | 0.89 | 0.66 | 56.99 | 46.78 | 71.00 |
| | 11 | 2.94 | 1.53 | 0.76 | 54.23 | 54.98 | 71.67 |
| | 12 | 4.18 | 3.15 | / | 77.03 | 67.42 | / |
| | **Mean** | **3.41** | **1.33** | **0.81** | **63.13** | **51.95** | **73.06** |
| | **SD** | **1.29** | **0.82** | **0.14** | **11.41** | **7.63** | **3.17** |

| Precursor | Subject ID | Variability of cycle duration (a.u.) | | | Index of EMG Variability (a.u.) | | |
|---|---|---|---|---|---|---|---|
| | | Birth | 3 months | Walking | Birth | 3 months | Walking |
| **Stepping** | 1 | 0.51 | 0.32 | 0.31 | 0.22 | 0.16 | 0.15 |
| | 4 | 0.19 | 0.24 | 0.20 | 0.20 | 0.17 | 0.15 |
| | 5 | 0.30 | 0.50 | 0.22 | 0.21 | 0.21 | 0.16 |
| | 8 | 0.59 | 0.44 | 0.09 | 0.30 | 0.15 | 0.09 |
| | 9 | 0.29 | 0.44 | 0.08 | 0.21 | 0.14 | 0.15 |
| | 10 | 0.37 | 0.37 | 0.11 | 0.30 | 0.11 | 0.10 |
| | **Mean** | **0.38** | **0.39** | **0.17** | **0.24** | **0.16** | **0.13** |
| | **SD** | **0.15** | **0.09** | **0.09** | **0.05** | **0.03** | **0.03** |
| **Kicking** | 2 | 0.50 | 0.20 | 0.12 | 0.31 | 0.13 | 0.13 |
| | 3 | 0.48 | 0.52 | / | 0.26 | 0.14 | / |
| | 4 | 0.21 | 0.32 | 0.20 | 0.23 | 0.14 | 0.15 |
| | 5 | 0.48 | 0.58 | 0.22 | 0.14 | 0.20 | 0.16 |
| | 6 | 0.60 | 1.06 | 0.26 | 0.23 | 0.12 | 0.09 |
| | 7 | 0.15 | 0.70 | 0.14 | 0.12 | 0.11 | 0.10 |
| | 8 | 0.71 | 0.73 | 0.09 | 0.13 | 0.19 | 0.09 |
| | 11 | 0.46 | 0.51 | 0.09 | 0.20 | 0.15 | 0.09 |
| | 12 | 0.57 | 0.33 | / | 0.16 | 0.14 | / |
| | **Mean** | **0.46** | **0.55** | **0.16** | **0.20** | **0.15** | **0.12** |
| | **SD** | **0.18** | **0.26** | **0.07** | **0.07** | **0.03** | **0.03** |
