## Supplemental Table 3 for "Generating variability from motor primitives during infant locomotor development"

**Table S3.**
P-values associated with Wilcoxon tests.
P-values are considered significant for $p<0.05$ (dark grey) and a trend is considered between 0.05 and 0.1 (light grey)
E1: birth / E2: 3 months old / E3: walking onset

| | | Stepping | | | Kicking | | |
|---|---|---|---|---|---|---|---|
| | | E1 - E2 | E2 - E3 | E1 - E3 | E1 - E2 | E2 - E3 | E1 - E3 |
| Cycle duration | | 0.031 | 0.031 | 0.031 | 0.004 | 0.110 | 0.016 |
| Proportion of phases | | 0.844 | 0.219 | 0.562 | 0.074 | 0.016 | 0.078 |
| Variability of cycle duration | | 0.687 | 0.031 | 0.062 | 0.301 | 0.016 | 0.016 |
| IEV | | 0.063 | 0.063 | 0.031 | 0.129 | 0.047 | 0.031 |
| VAF (Goodness of fit) for 4 modules | | 0.156 | 0.094 | 0.063 | 0.734 | 0.016 | 0.047 |
| For each individual's number of module | IRV | 0.156 | 0.031 | 0.031 | 0.734 | 0.016 | 0.016 |
| | IRS | 0.563 | 0.156 | 0.563 | 0.652 | 0.652 | 0.047 |
| | SMAI | 0.438 | 0.031 | 0.063 | 0.250 | 0.016 | 0.016 |
| | STAI | 0.563 | 0.031 | 0.094 | 0.301 | 0.016 | 0.016 |
| For a fixed number of 4 modules (methodological verification) | IRV | 0.063 | 0.063 | 0.031 | 0.039 | 0.016 | 0.016 |
| | IRS | 0.219 | 0.031 | 0.313 | 0.570 | 0.219 | 0.047 |
| | SMAI | 0.438 | 0.031 | 0.031 | 0.301 | 0.016 | 0.016 |
| | STAI | 0.688 | 0.031 | 0.031 | 0.301 | 0.031 | 0.016 |
