## Supplemental Table 4 for "Generating variability from motor primitives during infant locomotor development"

**Table S4.**

Individual data regarding the dimensionality of the signals (VAF for a modeling of 4 modules and number of modules to reach the threshold VAF).

The goodness of fit is arbitrarily considered sufficient above 75% (grey cells).

| Precursor | Subject ID | VAF (Goodness of fit) for 4 modules | | | Number of modules to reach VAF>75% | | |
|---|---|---|---|---|---|---|---|
| | | Birth | 3 months | Walking | Birth | 3 months | Walking |
| **Stepping** | 1 | 0.913 | 0.768 | 0.591 | 3 | 4 | 7 |
| | 4 | 0.725 | 0.772 | 0.637 | 5 | 4 | 6 |
| | 5 | 0.826 | 0.819 | 0.688 | 3 | 3 | 5 |
| | 8 | 0.820 | 0.769 | 0.754 | 4 | 4 | 4 |
| | 9 | 0.883 | 0.768 | 0.655 | 3 | 4 | 6 |
| | 10 | 0.713 | 0.688 | 0.747 | 5 | 5 | 5 |
| | **Mean** | **0.81** | **0.76** | **0.68** | **3.83** | **4.00** | **5.50** |
| | **SD** | **0.08** | **0.04** | **0.06** | **0.98** | **0.63** | **1.05** |
| **Kicking** | 2 | 0.728 | 0.842 | 0.749 | 5 | 3 | 5 |
| | 3 | 0.728 | 0.789 | / | 5 | 4 | / |
| | 4 | 0.769 | 0.704 | 0.637 | 4 | 5 | 6 |
| | 5 | 0.804 | 0.759 | 0.688 | 4 | 4 | 5 |
| | 6 | 0.845 | 0.891 | 0.636 | 4 | 3 | 6 |
| | 7 | 0.878 | 0.866 | 0.550 | 3 | 3 | 7 |
| | 8 | 0.767 | 0.799 | 0.754 | 4 | 4 | 4 |
| | 11 | 0.758 | 0.821 | 0.614 | 4 | 3 | 7 |
| | 12 | 0.795 | 0.710 | / | 4 | 5 | / |
| | **Mean** | **0.79** | **0.80** | **0.66** | **4.11** | **3.78** | **5.71** |
| | **SD** | **0.05** | **0.07** | **0.07** | **0.60** | **0.83** | **1.11** |
