## Supplemental Table 5 for "Generating variability from motor primitives during infant locomotor development"

**Table S5.**
Individual data for other indexes (IRV, IRS, SMAI, STAI).

| Precursor | Subject ID | IRV | | | IRS | | |
|---|---|---|---|---|---|---|---|
| | | Birth | 3 months | Walking | Birth | 3 months | Walking |
| **Stepping** | 1 | 34.82 | 15.06 | 4.12 | 0.43 | 0.37 | 0.54 |
| | 4 | 12.69 | 14.35 | 5.19 | 0.38 | 0.46 | 0.51 |
| | 5 | 25.36 | 28.79 | 7.16 | 0.48 | 0.37 | 0.40 |
| | 8 | 25.61 | 14.93 | 5.23 | 0.47 | 0.37 | 0.54 |
| | 9 | 27.86 | 11.07 | 5.73 | 0.51 | 0.44 | 0.49 |
| | 10 | 19.54 | 7.58 | 5.55 | 0.44 | 0.39 | 0.47 |
| | **Mean** | **24.31** | **15.30** | **5.50** | **0.45** | **0.40** | **0.49** |
| | **SD** | **7.53** | **7.22** | **0.99** | **0.04** | **0.04** | **0.05** |
| **Kicking** | 2 | 17.45 | 25.54 | 6.68 | 0.43 | 0.34 | 0.49 |
| | 3 | 18.82 | 13.60 | / | 0.38 | 0.45 | / |
| | 4 | 22.54 | 8.98 | 5.19 | 0.40 | 0.42 | 0.51 |
| | 5 | 17.22 | 16.29 | 7.16 | 0.35 | 0.38 | 0.40 |
| | 6 | 23.71 | 16.75 | 3.45 | 0.38 | 0.38 | 0.43 |
| | 7 | 17.99 | 18.34 | 2.53 | 0.35 | 0.49 | 0.37 |
| | 8 | 11.89 | 20.97 | 5.23 | 0.33 | 0.43 | 0.54 |
| | 11 | 19.15 | 22.57 | 2.90 | 0.50 | 0.44 | 0.46 |
| | 12 | 13.54 | 7.45 | / | 0.50 | 0.43 | / |
| | **Mean** | **18.04** | **16.72** | **4.74** | **0.40** | **0.42** | **0.46** |
| | **SD** | **3.76** | **6.01** | **1.83** | **0.06** | **0.04** | **0.06** |

| Precursor | Subject ID | SMAI | | | STAI | | |
|---|---|---|---|---|---|---|---|
| | | Birth | 3 months | Walking | Birth | 3 months | Walking |
| **Stepping** | 1 | 0.35 | 0.73 | 0.80 | 0.30 | 0.38 | 0.45 |
| | 4 | 0.69 | 0.78 | 0.84 | 0.40 | 0.35 | 0.44 |
| | 5 | 0.81 | 0.76 | 0.80 | 0.42 | 0.42 | 0.49 |
| | 8 | 0.77 | 0.75 | 0.85 | 0.45 | 0.40 | 0.45 |
| | 9 | 0.77 | 0.76 | 0.86 | 0.42 | 0.39 | 0.44 |
| | 10 | 0.72 | 0.83 | 0.92 | 0.41 | 0.38 | 0.41 |
| | **Mean** | **0.68** | **0.77** | **0.85** | **0.40** | **0.39** | **0.45** |
| | **SD** | **0.17** | **0.04** | **0.04** | **0.05** | **0.02** | **0.03** |
| **Kicking** | 2 | 0.78 | 0.75 | 0.79 | 0.42 | 0.37 | 0.47 |
| | 3 | 0.70 | 0.68 | / | 0.40 | 0.35 | / |
| | 4 | 0.67 | 0.71 | 0.84 | 0.38 | 0.44 | 0.44 |
| | 5 | 0.72 | 0.74 | 0.80 | 0.42 | 0.40 | 0.49 |
| | 6 | 0.63 | 0.84 | 0.85 | 0.38 | 0.42 | 0.47 |
| | 7 | 0.68 | 0.68 | 0.88 | 0.39 | 0.35 | 0.47 |
| | 8 | 0.70 | 0.69 | 0.85 | 0.40 | 0.34 | 0.45 |
| | 11 | 0.68 | 0.77 | 0.78 | 0.41 | 0.37 | 0.47 |
| | 12 | 0.72 | 0.74 | / | 0.44 | 0.44 | / |
| | **Mean** | **0.70** | **0.73** | **0.83** | **0.40** | **0.39** | **0.47** |
| | **SD** | **0.04** | **0.05** | **0.04** | **0.02** | **0.04** | **0.02** |
