## Supplemental Table 6 for "Generating variability from motor primitives during infant locomotor development"

**Table S6.**

Individual data of 12 adults from Hinnekens et al. (2020) to plot Figure 1 as well as adult landmarks on Figures 4 and 5. Note that those data are displayed for illustration: even though we retrieved raw data to apply the same filtering and normalization as in the rest of the paper, data are not directly comparable since adult cycles were defined as step cycles (with a stance phase and a swing phase) whereas infant/toddler data were cut off as flexion and extension cycles (with an extension phase and a flexion phase).

| **Adult** | **IEV** | **VAF** | **Module number to reach VAF>75 %** | **IRV** | **IRS** | **SMAI** | **STAI** |
|---|---|---|---|---|---|---|---|
| 1 | 0.10 | 0.63 | 6 | 2.28 | 0.62 | 0.69 | 0.39 |
| 2 | 0.08 | 0.73 | 5 | 2.45 | 0.61 | 0.65 | 0.41 |
| 3 | 0.11 | 0.67 | 5 | 3.66 | 0.63 | 0.65 | 0.41 |
| 4 | 0.10 | 0.78 | 4 | 4.71 | 0.59 | 0.67 | 0.38 |
| 5 | 0.09 | 0.78 | 4 | 4.42 | 0.58 | 0.63 | 0.45 |
| 6 | 0.09 | 0.72 | 5 | 2.20 | 0.64 | 0.65 | 0.42 |
| 7 | 0.11 | 0.71 | 5 | 4.41 | 0.60 | 0.68 | 0.40 |
| 8 | 0.10 | 0.74 | 5 | 3.81 | 0.58 | 0.61 | 0.46 |
| 9 | 0.09 | 0.73 | 5 | 3.17 | 0.63 | 0.62 | 0.46 |
| 10 | 0.10 | 0.82 | 4 | 4.46 | 0.59 | 0.64 | 0.41 |
| 11 | 0.08 | 0.69 | 5 | 2.50 | 0.62 | 0.62 | 0.39 |
| 12 | 0.10 | 0.72 | 5 | 3.36 | 0.61 | 0.64 | 0.42 |
| **Landmark (mean)** | **0.10** | **0.73** | **4.83** | **3.45** | **0.61** | **0.65** | **0.42** |
| **Standard deviation** | **0.01** | **0.05** | **0.58** | **0.93** | **0.02** | **0.02** | **0.03** |
